# Fitness effects of breeding strategy: implications for life history trait evolution and mouse husbandry

**DOI:** 10.1101/2023.02.13.526889

**Authors:** Alexis Garretson, Beth L. Dumont

## Abstract

Reproductive tactics can profoundly influence population reproductive success, but paradoxically, breeding strategy and female reproductive care often vary across a population. The causes and fitness impacts of this variation are not well understood. Using breeding records from the Collaborative Cross mouse population, we evaluate the effects of breeding configuration on reproductive output. Overall, we find that communal breeding in trios leads to higher output and that both trio-breeding and overlapping litters are associated with increased neonatal survival. However, we find significant strain-level variation in optimal breeding strategy and show that the tradeoff between strategies is weakly heritable. We further find that strain reproductive condition influences the ability to support multiple litters and alters the related evolutionary tradeoffs of communal breeding. Together, these findings underscore the role of genetics in regulating alternative reproductive tactics in house mice and emphasize the need to adopt animal husbandry practices tailored to strain backgrounds.

## Introduction

A critical goal in evolutionary biology is to understand how phenotypic variation is maintained in populations.^1,2^ Population-level variations in reproduction-related life-history traits among individuals of the same sex are particularly perplexing, as such traits can lead to variance in reproductive success,^2,3^ and should be subjected to intense selection toward the most optimal strategy. However, alternative reproductive tactics (ARTs) are observed in a variety of taxa, including fish, birds, reptiles, amphibians, insects, and mammals.^4^

One such successful and pervasive reproductive tactic observed across animal taxa is alloparenting, where a conspecific individual other than a genetic parent provides parental care to an offspring. Examples of alloparenting have been documented in many species, including more than 120 mammals across most orders.^5^ In humans, alloparental care is universal,^6^ is linked to increased fecundity and childhood survival,^7,8^ and has likely played crucial roles in human cultural and cognitive evolution.^6,9^ However, despite its prevalence and significance, the genetic and neurobiological causes and correlates of alloparenting remain poorly understood.^10^

In house mice (*Mus musculus*), alloparenting is observed in both the laboratory and in wild mouse populations and may include both female communal care and communal nursing.^11–13^ In the wild, house mouse social structure is variable but typically includes multiple breeding females per group along with a single dominant male and, often, several male or female non-breeding subordinate individuals.^14,15^

Communal nesting and nursing in female mice have a variety of costs and benefits to individual females and mouse populations. On the one hand, communal care can lead to the exploitation of secondary females if litter sizes or energetic investment in nursing are not equal, reducing the relative fitness of the secondary female.^16,17^ Additionally, communal breeding can be associated with higher rates of within-nest infanticide.^18,19^ These costs of communal breeding may explain why free-ranging mice reportedly nurse communally and pool litters only when a close female relative is available.^20^ On the other hand, despite these costs, communal breeding has been demonstrated to increase the lifetime reproductive success of both females and increase pup survival ^12,21^. However, these benefits may depend on underlying individual-level differences in reproductive success and correlated phenotypes. For example, in many species, reproductive fitness traits are a function of female body mass and age;^22,23^ increased body mass leads to higher milk production and a higher probability of dominance over smaller females.^17,24,25^ As a result, females with greater body mass raise a higher proportion of solitary litters.^23^ These complex cost-benefit relationships suggest that communal and solitary care may be differentially favored under different environmental or population conditions, and when the health and reproductive quality of individual females varies.

Because female mice can alternate between solitary and communal breeding during their lifetime, breeding tactics in female mice are believed to be phenotypically plastic rather than solely genetically controlled.^23,26^ However, even between congeneric mouse species, there is significant variation in breeding strategies.^27^ For example, *Mus musculus* is considered polygynous or promiscuous while *Mus spicilegus* is monogamous, suggesting that there may be some level of genetic control of breeding phenotypes and social group behavior.^14,28,29^ Determining the relative roles of genetics and environment to the magnitude of the fitness tradeoffs between solitary and communal breeding could offer new insights into the evolutionary stability of these alternative reproductive strategies.

Beyond its broad relevance for evolutionary biology, understanding to what extent breeding behavior alters reproductive success is of critical importance in research colonies of laboratory mice. House mice are one of the most widely used animal models in biological research, partly because they are relatively easy and inexpensive to maintain in a laboratory setting and are prolific breeders.^30,31^ However, constraints on laboratory mouse housing and husbandry and consideration of the limited space in animal facilities have led to the utilization of multiple breeding designs to maximize reproductive output or experimental success.^32,33^ Two common strategies for laboratory mouse breeding are continuous trio and continuous pair breeding. Trio mating, which is established by co-housing two female mice with a single male mouse, is a common rapid research colony expansion strategy because it may increase reproductive performance with lower demand on space and is comparable to communal care in free-living populations.^32,34,35^ In the laboratory, trio breeding designs have been associated with increased pup growth weight, larger litter sizes at wean, and higher body weights as adults, with no adverse impacts on pup welfare.^11,33,34,36,37^ However, other investigations have found that pairs outperform trios in laboratory settings because of reproductive suppression of one breeding female in the trio,^19^ high pre-weaning lethality in trios with overlapping litters,^38^ and within-nest infanticide.^12,23^

Previous studies investigating the effects of breeding configuration on reproductive success have focused on only one or a small number of well-characterized inbred laboratory mouse strains. In particular, comparative studies of wild and laboratory house mice suggested that wild mice may have more extreme copulatory behavior than laboratory mice, indicating that some genetic elements influencing reproductive behavior may not be found in standard laboratory strains.^39^ Moreover, wild mouse studies often employ small sample sizes and data collection regimes that are not strictly comparable to those in a production breeding facility. Notably, the controlled setting of a production-scale breeding environment enables facile exploration of large-scale, long-term breeding strategy tradeoffs.

In this study, we use a dataset of 4,540 crosses from 53 genetically diverse Collaborative Cross (CC) inbred mouse strains reared in a single breeding facility to investigate the impacts of breeding configuration on reproductive success. The CC are a recombinant inbred panel of mice developed from eight diverse founder strains, with each CC strain genome representing a unique genetic mosaic of the eight founder strains. In particular, we (1) investigate the magnitude of the fitness tradeoff between pair and trio breeding configurations, (2) determine the influence of breeding configuration and overlapping litters on litter survival rates, (3) catalog the variability in the fitness tradeoffs across strains from genetically distinct backgrounds and estimate the proportion of variance due to genetic differences, and (4) determine to what extent the fitness consequences result from female and male reproductive conditions. Importantly, by profiling a large panel of genetically diverse strains, our study design allows the first rigorous analysis of how genetic background modulates reproductive success as a function of breeding configuration.

## Results

### Reproductive Traits vary as a function of Breeding Configuration across CC strains

We collated multiple reproduction-associated measures from laboratory breeding records for 53 CC strains (Figure 1). There are significant strain-level differences for all surveyed measures (one-way ANOVA P < 0.0001), establishing considerable variation in reproduction-related traits across this multiparent mapping population. In addition, breeding success metrics were generally highly correlated between trio and pair breeding configurations (Figure 1), with an average correlation coefficient of 0.75±0.17.

**Figure 1.**
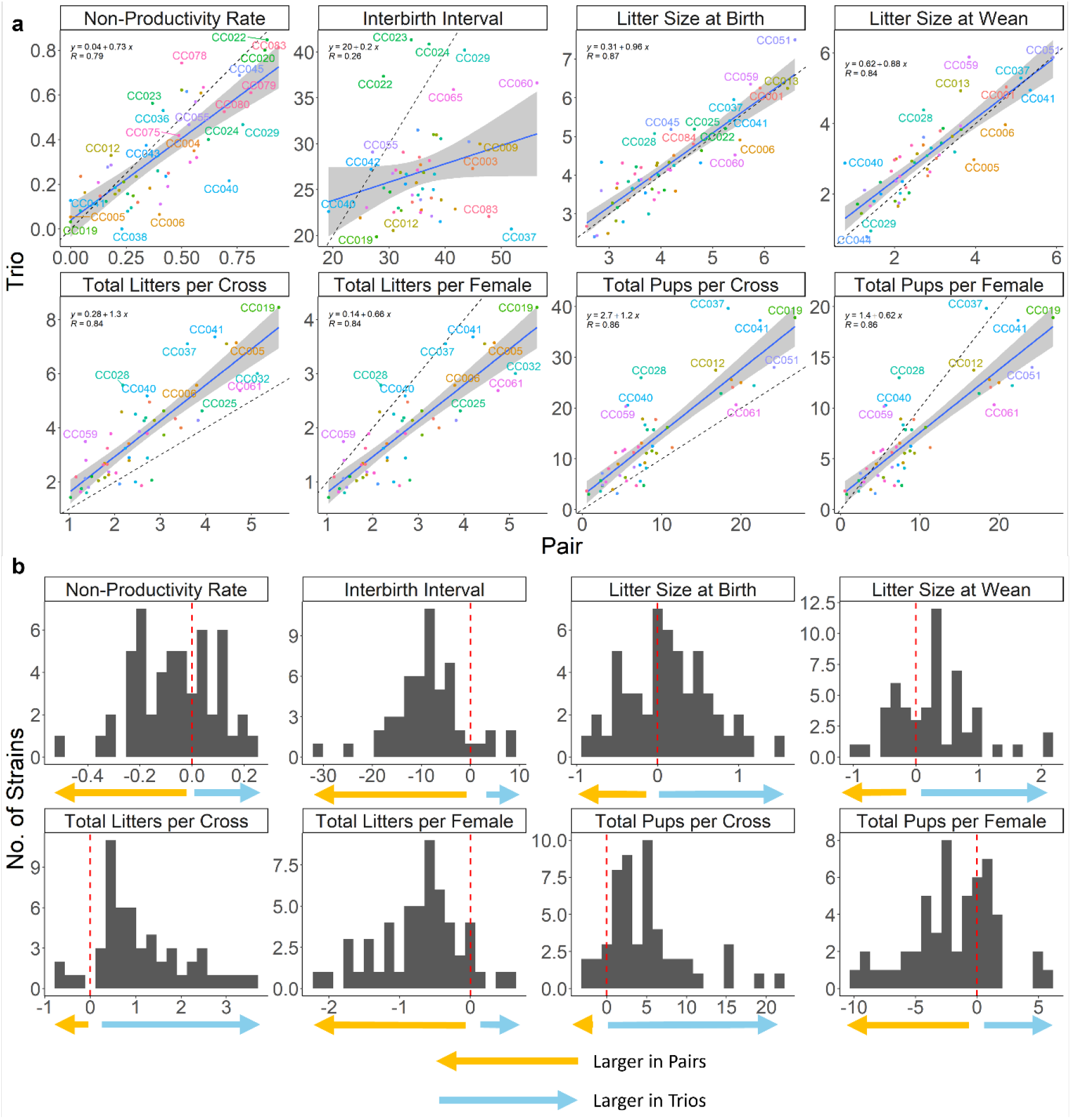
A) Scatter plots comparing the trio and pair values for each strain for each surveyed reproductive metric. Dotted lines show x=y, the blue line shows a fitted linear regression, and the grey-shaded area shows standard error. Equations of the lines and the correlation coefficients are presented in the upper left. Points are labeled with the corresponding Collaborative Cross strain. B) Histograms of trio-pair divergence values for each strain for each surveyed reproductive metric. Dashed red line indicates the parity of trio and pair values. Arrows denote the portion of the graph where the assayed value was larger in pairs (orange) or trios (blue).

A common, implicit motivation for establishing breeding trios assumes that this mating configuration will increase reproductive output. Consistent with this expectation, we observed an increased frequency of litters (F(1,4434)=453.51, P<0.0001), decreased interbirth interval (F(1,10007), 278.55, P<0.0001), larger litter sizes at birth and wean (F(1,13010)=974, P<0.0001, F(1,13010)=974, P<0.0001), increased number of pups per cross (F(1, 4452)=29.68, P<0.001), and a nearly doubled probability of cross productivity (z=-5.46, P<0.0001; Figure 2) in trio compared to pair matings across the entire CC population. In aggregate, the frequency of litters weaned from trio mating units was not double that of pairs, with trios weaning only 1.68 times as many litters compared to pairs (3.93±3.14 versus 2.34±1.89). However, the total number of pups was more than doubled in trios (15.49±15.30) relative to pairs (7.02±8.93, 1.21 fold change, F(1, 4452)=29.68, P<0.001), reflecting the increase in average litter size at wean in trio compared to pair breeding configurations.

**Figure 2.**
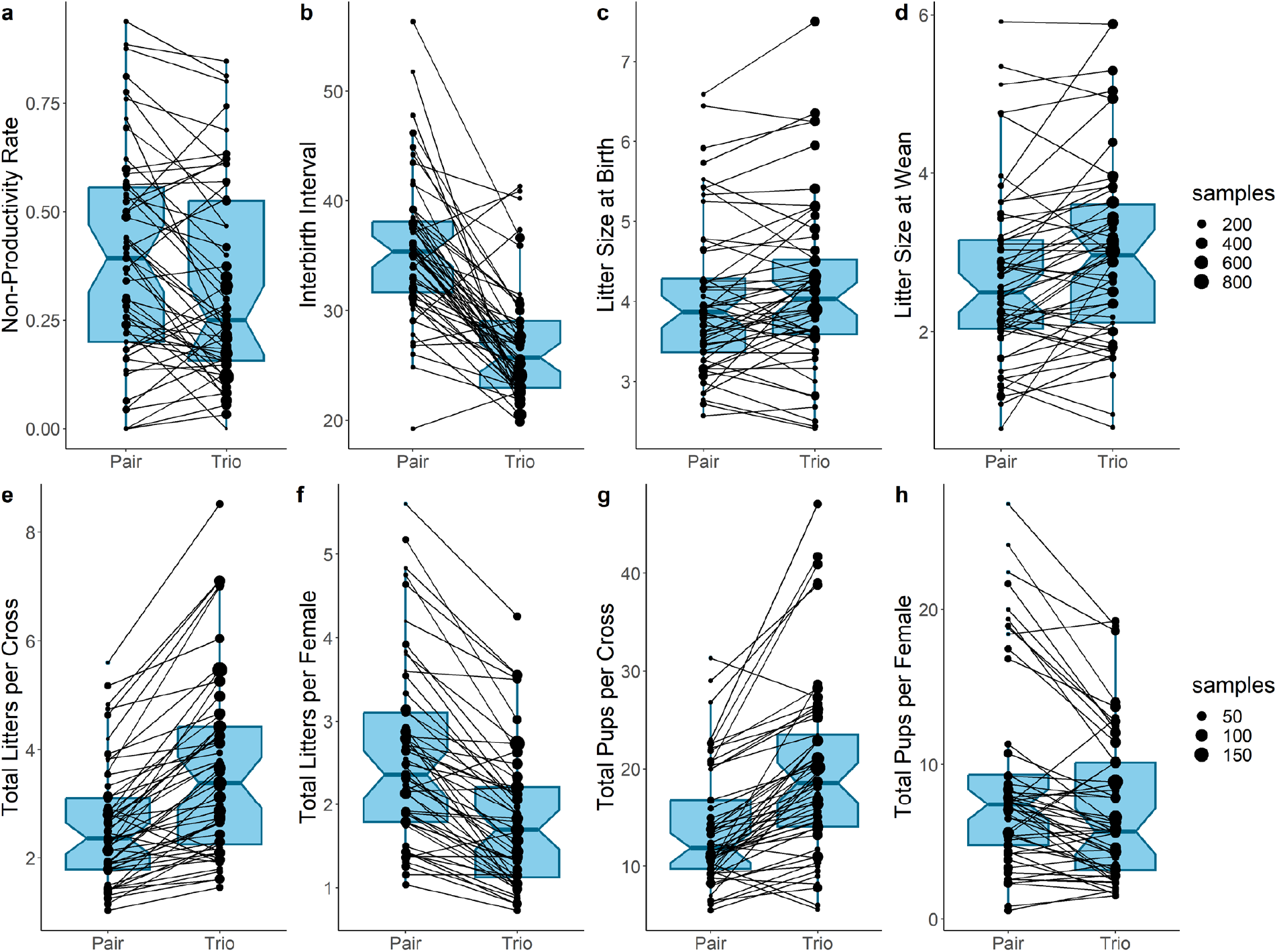
Boxplots comparing the strain-level values for each surveyed metric of breeding performance for breeding mouse trios and pairs: A) non-productivity rate, B) interbirth interval, C) litter size at birth, D) litter size at wean, E) total number of litters per cross, F) total number of litters per female, G) total number of pups per cross, and H) total number of pups per female. Lines connect the trio and pair values for each strain, and points are sized by the number of samples (i.e., crosses or litters) in that comparison. All comparisons are significant at *P* < 0.05.

For individual strains, the total number of pups born per cross ranged from 0.61 (CC023) to 6.56 (CC083) times greater in trios than for pairs, and the total number of litters per cross ranged from 0.72 (CC078) to 2.57 (CC059) times larger in trios than pairs. Thus, while trio breeding configurations were associated with increased overall breeding performance and productivity compared to pair mating designs across the CC population, reproductive performance metrics for some individual strains actually decrease under a trio configuration (Figure 2). In addition, two-way ANOVA revealed a significant interaction between the effects of breeding design and strain for several reproduction phenotypes, including total litters per cross (F(52, 4366)=2.086, P<0.0001), total pups per cross (F(52, 4452)=317, *P*<0.0001), litter sizes at birth (F(52, 13010)=1.46, *P*=0.017), and litter sizes at wean (F(52, 13834)=13.8, P<0.0001).

These findings reveal strain-to-strain variation in reproductive performance under different breeding paradigms and suggest that the optimal breeding strategy for maximizing reproductive output is strain-and genotype-dependent.

### Significant Strain-Level Variation and Heritability in Reproductive Performance

The difference in reproductive performance between trio and pair breeding units provides an estimate of the relative fitness gain (or loss) associated with these two ARTs. Each reproductive metric exhibits at least one strain for which trio breeding units outperform pair mating units, and at least one strain with pair breeding units outperforming trio matings (Figure 1). For several traits, including the non-productivity rate and the litter size at birth, there is a near-equal number of strains that outperform in trio and pair configurations.

Trio-pair differentials are strongly correlated with many reproductive traits (Figure 3). For example, strains with larger average litter sizes have larger trio litter sizes at birth (r=0.29, *P*=0.035) and wean (r=0.32, *P*=0.017), and more litters per cross relative to pairs (r=0.31, *P*=0.025; Figure 3B). Additionally, shorter average interbirth intervals and lower ages at first litters were associated with more litters per cross relative to pairs, higher survival rates relative to pairs, and lower whole litter loss rates in trios relative to pairs. Finally, earlier ages at first litter were associated with shorter interbirth intervals in trios relative to pairs.

**Figure 3.**
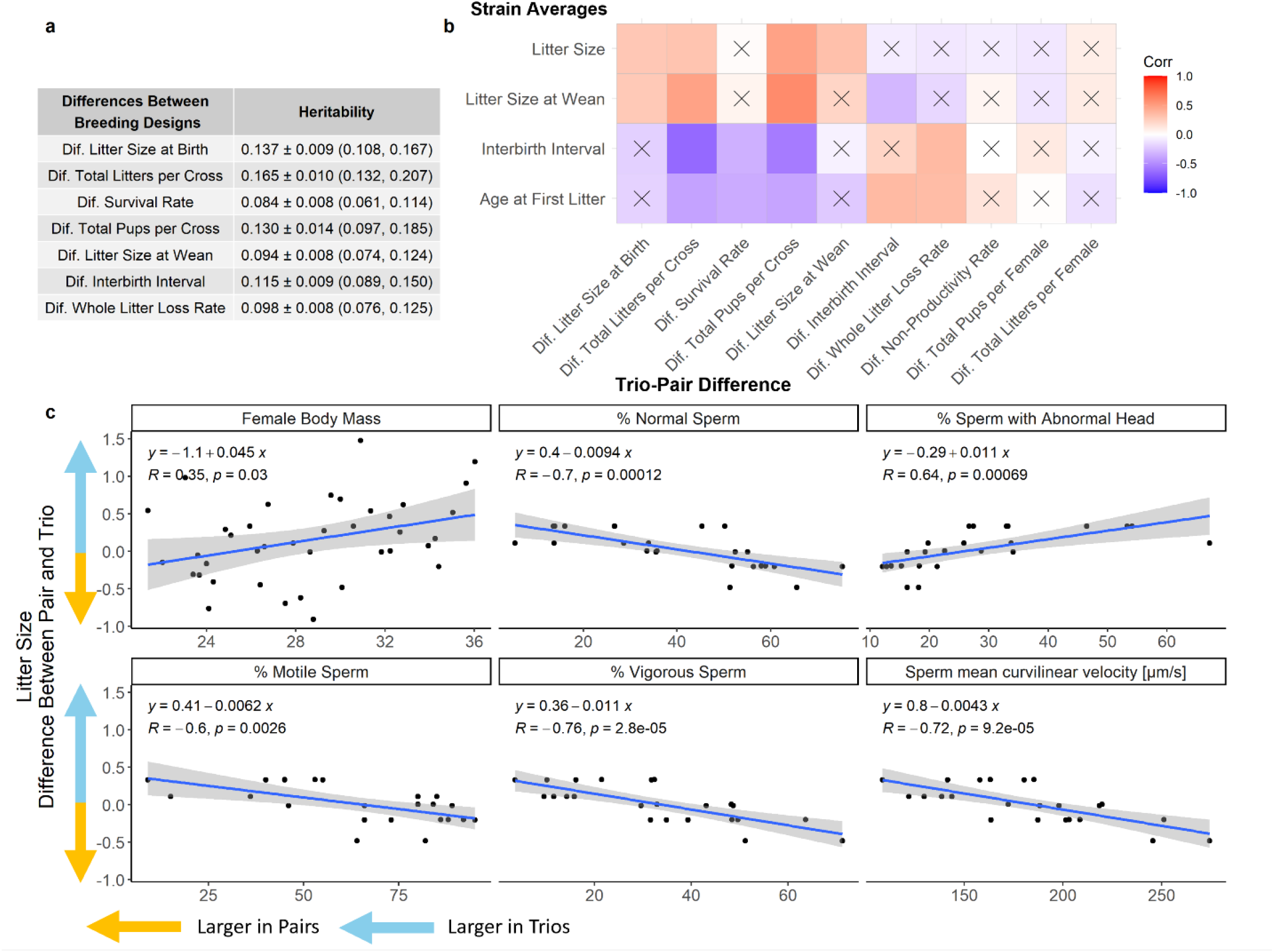
A) Estimated heritabilities for the trio-pair divergence values for each reproductive metric. B) Heatmap showing the correlation between the overall strain average value for a reproductive fitness trait and the trio-pair differences for the target traits. Boxes labeled with “X” correspond to non-significant correlations. C) Correlations between the difference in litter size between trios and pairs and both strain average female body mass and metrics of sperm condition. The blue line shows a fitted linear regression, with the grey-shaded area showing standard error. Equations of the lines and the correlation coefficients are presented in the upper left.

We used analysis of variance to estimate the heritability of the trio-pair differential for breeding performance metrics (Figure 3A). The heritability of these traits varied between 0.084 (trio-pair divergence in survival rates) to 0.165 (trio-pair divergence in total litters per cross), indicating that these trait differentials are weakly heritable. Heritability estimates for each trait, independently derived for trios and pair matings, are notably greater than the trait differentials (an average of 1.94 times greater) and are provided in (Supplementary Table 1).

We performed QTL mapping scans to localize genetic regions contributing to observed variation in the trio-pair differential for various reproductive performance metrics in the CC. We identify suggestive QTL peaks with LOD scores >6 (roughly corresponding to α= 0.1) for litters per cross, interbirth interval, and the probability of a non-productive cross on chrs 6, 1, and 5, respectively. However, none of these QTL is significant at *P*<0.05, and peaks were too broad to highlight individual putative candidates contributing to the observed variability (Supplementary Figure 1).

### Reproductive Trait Differential between trios and pairs is associated with body size and sperm quality

We next sought to determine whether the observed differences in breeding performance between trios and pairs could be explained, at least in part, by strain-level variance in measures of overall reproductive condition. Trait differences between breeding configurations were significantly correlated with the strain average female body mass (*P*=0.03, R=0.35), with strains characterized by larger adult female body masses producing larger litter sizes in trios relative to pairs (Figure 4C). In addition, trio-pair divergence in litter sizes was highly negatively correlated with several strain-level metrics of sperm quality, including velocity (*R*=-0.72, *P*<0.0001), proportion of sperm with normal morphology (*R*=-0.7, *P*=0.0001), proportion of motile sperm (*R*=-0.6, *P*=0.003), proportion of vigorous sperm (*R*=-0.76, *P*<0.001), and the proportion of sperm with abnormal heads (*R*=0.64, *P*=0.0007, Figure 4C). Thus, strains with higher maternal body weight or paternal sperm quality often experienced greater fitness gains when bred in a pair configuration rather than as trios (Figure 3C).

**Figure 4.**
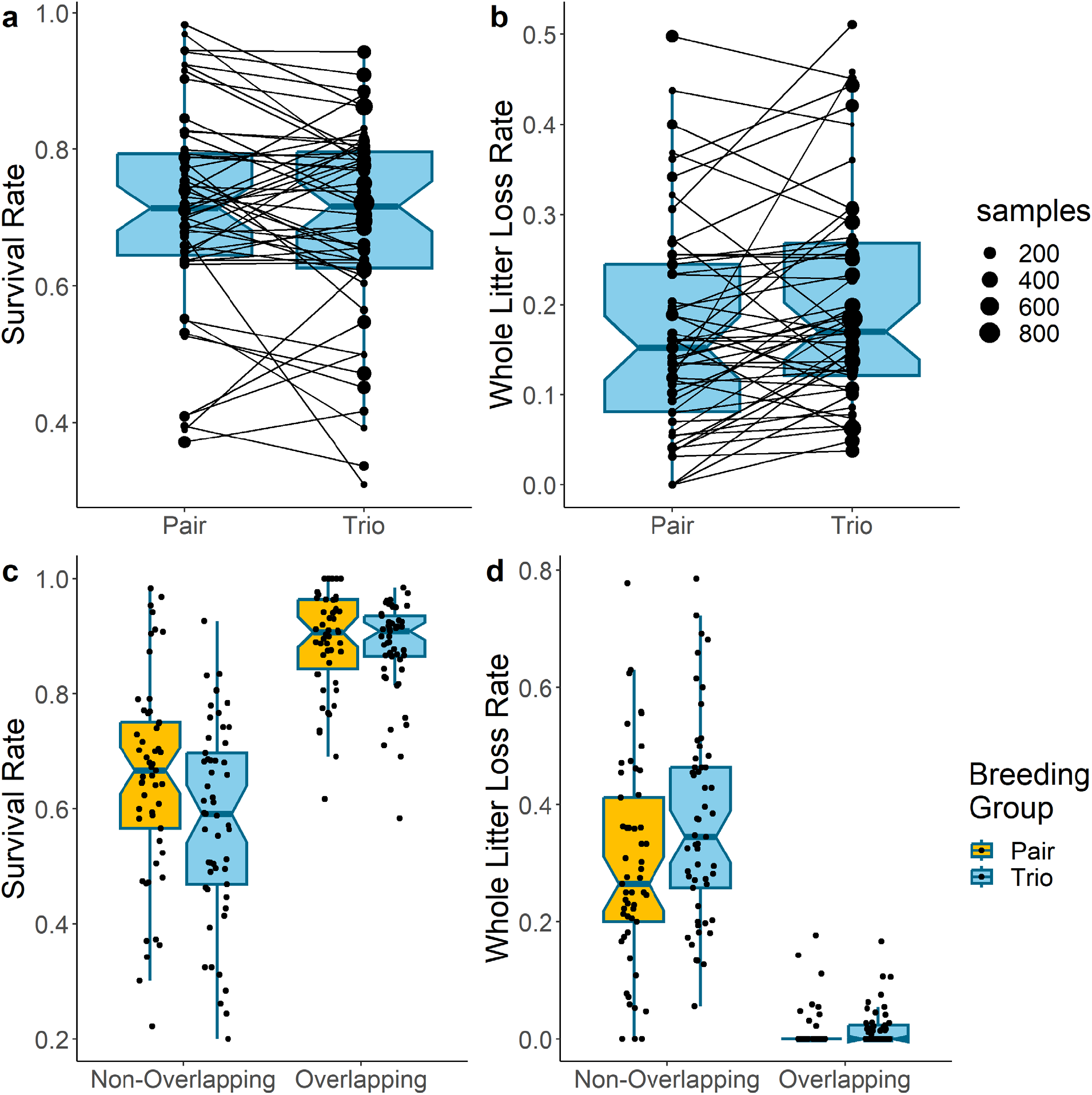
Boxplots comparing the strain level values for trios and pairs for A) the survival rate and B) the whole litter loss rate. Lines connect the trio and pair values for each strain, and points are sized by the number of samples in that comparison. All comparisons are significant at P < 0.05. Boxplots comparing the survival rate (C) and whole litter loss rate (D) between trio and pair breeding configurations stratified by the presence of an overlapping litter.

### Higher Pup Survival Rates are Associated with the Presence of Overlapping Litters

Laboratory mouse pup mortality is a significant economic and animal welfare concern, but its causes are poorly understood. We report a significant effect of breeding design on litter survival rates (F(1,12798)=15.13, *P*=0.0001, Figure 4A), with overall trio survival proportions (0.73±0.40) being significantly higher than pair survival proportions (0.69±0.41). However, like other reproductive traits, this general trend is not uniform across strains. Individual CC lines show differences in the magnitude and sign of the difference in pup survival between trio and pair configurations (two-way ANOVA modeling survival proportion as a function of breeding design and strain, F(52, 12798)=1.94, P<0.0001).

Remarkably, including predictors for overlapping litters and increased litter sizes, a linear mixed model yields lower survival rates for litters in trio breeding configurations (−0.069±0.008, t=-8.57, *P*<0.000). Thus, the association between litter survival and breeding configuration may be primarily mediated by the presence of littermates or overlapping litters. In both trios and pairs, the presence of an overlapping litter is associated with a significant increase in the survival rate (0.25±0.01, t=24.43, P<0.0001, Figure 2C), with survival rates increasing for each additional day that overlapping litters are co-housed (0.0029±0.0006, t=4.17, P<0.0001). In contrast, the relationship between litter survival and breeding configuration is not significantly modified by the number of overlapping siblings (*P*=0.48), although increasing litter sizes were associated with increased survival rates (0.052±0.001, t=36.71, P<0.0001). Further, we did not detect a significant effect of the proportion of a litter surviving to wean (*P*=0.93) or the number of previous litters (*P*=0.37) on pup survival rates.

Previous work suggests that reproductive behavior surrounding whole litter loss may be distinct from other forms of pup mortality.^40,41^ We observe a significant effect of breeding configuration on the whole litter loss rate, with pairs exhibiting lower rates of litter loss than trios (GLMM Estimate: 0.26±0.035, z=7.36, *P*<0.001, Figure 2B). As with the litter survival rate, the probability of losing a whole litter decreased with the presence of an overlapping litter (GLMM Estimate -2.79±0.09, z=-30.25, P<0.0001, Figure 2D), as well as with each additional overlapping day (GLMM Estimate -0.042±0.001, z=-4.44, P<0.0001). The whole litter loss rate was also decreased with each additional overlapping sibling (GLMM Estimate -0.037±0.006, z=-6.42, P<0.0001). The probability of losing a whole litter did increase with the increasing age of the dam (GLMM Estimate 0.003±0.0004, z=5.9, P<0.0001), but it did not change significantly with litter number (*P*=0.49).

## Discussion

### Differential fitness effects of communal and solitary breeding in laboratory mice

The adoption of both trio and pair mating configurations across the standardized CC strain production environment presents a natural opportunity to investigate how different breeding strategies impact reproductive fitness. We harnessed the unique strengths of this genetically diverse inbred mouse strain population to first show that forced communal and solitary breeding did not result in equal fitness at the colony level, as quantified by several metrics of reproductive output. Specifically, we find that trios outperform pairs at the colony level and that this performance gain is acquired by reducing the frequency of non-productive matings, increasing pup survival rates, and resulting in larger litter sizes at birth and weaning (Figure 2). Importantly, however, communal breeding does not maximize reproductive output at the level of individual females, as trio breeding configurations do not double the reproductive output of breeding pairs. Our work reveals the presence of crucial tradeoffs between individual and population-level reproductive success through communal breeding and suggests that there may be risks of exploitation or free-riding by one female in a communally breeding trio.^17,42^

### Strain-specific effects on reproductive output are related to reproductive condition

We demonstrate clear fitness gains associated with communal breeding at the whole-colony level. However, at the level of individual CC strains, we uncover widespread strain-specific effects on reproductive output and success under different breeding strategies, with strain-level variation in both the magnitude and direction of the fitness differential between pairs and trios (Figure 1). The variable conclusions reached in previous studies of communal breeding, with some seeing production benefits to trio breeding^11,33,34^ and others reporting significant costs^19,35,38^, may therefore be due to a restricted focus on a single strain or a small number of strains. We show that these strain effects are weakly heritable, suggesting that segregating genetic variation in the CC strains can influence the relative efficacy of different reproductive tactics. The genetic identity of the loci driving these strain-dependent trait differentials remains unknown. While the CC was initially intended to serve as a genetically diverse mapping population, too few strains are currently available to enable well-powered mapping studies.^43^ Our QTL mapping results did not find significant loci contributing to variability in trio-pair difference in reproductive output, but we do observe some suggestive QTL peaks with LOD scores >6. Future studies on larger mouse panels may offer the increased power necessary to reveal putative candidates contributing to the genetic architecture of these traits.

We observed a positive relationship between female body size and the trio-pair difference in litter size (Figure 4C). While earlier work has reported that free-living, heavier female mice are more likely to rear their litters solitarily,^23^ our findings uncover the opposite trend in captive lab mouse populations. Using body size as an overall proxy for reproductive condition and available energy stores for reproduction, we find that high-condition CC females have higher reproductive success under communal breeding designs. These contradictory findings may owe to differences in the genetic background of outbred wild mice and the inbred strains profiled here but could also be mediated by environmental differences between the laboratory and wild environment.

We also report that strain reproductive condition is associated with the tradeoffs from benefits from communal breeding. Across the CC population, trio matings are more than 15% less likely to be non-productive than pair matings, but this effect is most substantial in strains with low overall fecundity. Additionally, strains with defective sperm morphology or motility reap greater reproductive gains from breeding in a trio, as opposed to a pair mating configuration (Figure 4C). Thus, many CC strains with low reproductive success maximize their reproductive potential under trio breeding designs. Future investigations are needed to determine whether this trend extends to non-CC mouse strains. As many inbred mouse models suffer from poor breeding performance, understanding the generality of this trend would immensely benefit colony management strategies and animal breeding programs.

### Overlapping litters are linked to higher pup survival rates

Laboratory mouse pup mortality is a significant economic and animal welfare concern, but its causes are diverse, ranging from environmental factors like temperature or nesting material availability to life history features such as dam age, litter size, and the level of parental care.^44,45^ While overlapping litters arise in both trio and pair breeding configurations, the phenomenon is more frequent in trios and significantly contributes to variation in litter survival rates.^38,45^ For example, previous work studying only trio-bred C57BL/6 animals showed a large increase in the whole litter loss rate of overlapping litters and a 2-7% increase in the probability of pup mortality.^38^

Conversely, in both pair and trio breeding configurations, we find that overlapping litters are associated with higher survival rates (Figure 4). This effect was very strong at the colony level but did vary across strains, suggesting that strain genetic background modulates the relative benefit of different breeding designs. In particular, the increased prevalence of overlapping litters appears to contribute to our finding that trios significantly increase survival rates (Figure 4). We speculate that communally bred females may be able to absorb the higher food intake costs of overlapping litters by communal nursing, effectively distributing the energy intake burden across multiple females.^12,46^ Our finding that larger average female body weights were associated with more successful communal breeding lends additional support to this interpretation (Figure 3C).

We also show that the increasing age of the overlapping litter (Figure 4) is associated with increased survival. This result raises the possibility that juvenile mice provide additional alloparenting benefits to younger litters.^47^ Underlying strain variance due to genetic background in the tendency for communal nursing, juvenile parental behavior, and sibling competition may further explain the appreciable variation in the survival rates of pups across strains and breeding configuration, as well as the spectrum of effects of communal breeding and overlapping litters observed across the Collaborative Cross strains.

The presence of overlapping litters in trios may facilitate higher levels of alloparenting from the accessory female by taking advantage of reproductive synchrony and facilitating indiscriminate communal care. However, we acknowledge that a limitation of this study is that pup data were only collected at the level of breeding units and are not available for individual females. Thus, we cannot directly assess potential reproductive suppression in either female in a trio. However, as most of the overlapping litters were born within ten days of the previous litter (60.2%), they are extremely unlikely to be born to a common dam. Further, in the vast majority of trios included in this study, both breeding females were littermates. Previous work has demonstrated that familiarity and genetic relatedness are critical components in the success of communal breeding in mice^48,49^ and humans.^50^ Future studies investigating the costs and benefits of different breeding strategies in the presence of overlapping litters of known parentage and using females with variable genetic relatedness and multiple genetic backgrounds could provide additional insight into the mechanisms controlling alternative reproductive tactics and reproductive fitness in house mice.

## Conclusions

By using the CC panel of genetically inbred mice assigned to different, ethologically relevant breeding configurations, we document significant strain variability in the fitness impacts of communal vs. solitary care, even under a fixed environment. Furthermore, we find that the reproductive fitness effects of these alternative breeding strategies are weakly heritable, establishing a genetic role for intraspecific variation in life history traits. Taken together, our study demonstrates the broad utility of the CC mouse panel for reproductive life-history investigations. Additionally, our findings provide insight into the extent of genetic control in the fitness tradeoff for alloparenting, a foundational trait in the study of human evolution^6^ and a key determinant of child survival rates across diverse human societies.^7,8^ More significantly, our work rigorously addresses a fundamental aim of evolutionary biology and population ecology by providing a new, mechanistic understanding of how variation in reproductive life-history strategies leads to differences in fitness.

## Methods

### Mice and housing

We used breeding records from Collaborative Cross mouse strains maintained at The Jackson Laboratory from 2016 to 2021, a subset of which were previously reported.^51^ The Collaborative Cross (CC) is a multiparent, recombinant inbred strain panel developed from eight founder strains, including three wild-derived strains.^52,53^ These eight founder strains are A/J, C57BL/6J, 129S1/SvImJ, NOD/ShiLtJ, NZO/HlLtJ, CAST/EiJ, PWK/PhJ, and WSB/EiJ. Each of the ∼70 extant CC strains was created via several generations of organized crossing, followed by at least 20 generations of brother-sister inbreeding to produce a reproducible genetic patchwork of the eight founder strains. The CC founder strains derive from three primary house mouse subspecies and capture nearly 90% of the total genetic variation and diversity observed in *Mus musculus*.^54–56^ Because of the inclusion of three wild-derived founders, these lines display considerable trait variation that is not found in common inbred strains that have been bred for ease of handling in the laboratory environment.^57,58^ We limited the dataset to those strains with at least five crosses and litters as well as at least ten total pups weaned in each breeding design (pairs and trios). The resulting retrospective mouse breeding data analyzed include 54,958 pups across 13,116 litters from 4,540 crosses from 53 CC strains.

### Statistical Methods

We retrieved breeding data from cage cards, including the strain, litter size, birth dates, parent identity, parent birth dates, and breeding configuration. Reproductive performance metrics analyzed include litter sizes at birth, the number of pups weaned per litter (from which we derived litter-level survival rates and the probability of whole litter loss), and the number of days between litters (interbirth interval). We also determined whether the cross was productive (i.e., yielded at least one live-born pup) and the total number of litters and pups per productive cross. This latter quantity was used to derive the total number of litters and pups per female.

For each reproductive success metric and strain, we stratified the data by breeding configuration and performed a simple linear regression of the mating pair value against the trio value. We next calculated the difference between the average trio and the pair-level reproductive trait values for each strain. Because these breeding metrics had only one value per strain, to calculate the heritability of the differences between solitary and communal breeding, we compared the difference between randomly sampled average trio and pair values from each cross for each metric to create a sample of 100 divergence values per strain. We then estimated the broad-sense heritability using the interclass correlation:

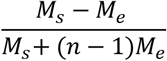

where *n* is the sample size, *M*_*s*_ is the strain mean square, and *M*_*s*_ is the residual mean square.^59^ We repeated this random sampling and heritability calculations 1,000 times per trait.

We carried out QTL mapping for each reproductive fitness trait using the linear mixed model approach implemented in r/qtl2 package.^60^ Mapping was performed using the CC presets, with non-random relatedness among strains accounted for by a kinship matrix tabulated using the leave-one-chromosome-out method. Significance was assessed by 1000 permutations of the empirical data.^61^

To evaluate the impact of breeding design and strain background on metrics of reproductive fitness, we independently modeled litter size, weaned litter size, survival rate, the interbirth interval for each litter, the total number of litters per productive cross, and the total number of pups per productive cross as functions of breeding configuration (pair or trio), strain identity, and the interaction between breeding design and strain using ANOVA. Strain and breeding configurations were treated as fixed factors. Tukey’s post hoc HSD test was used to find the differences between breeding configurations. Finally, we modeled the probability that a cross was unproductive as a general linear mixed model with a binomial distribution as a function of the breeding group and the age of the dam at the mating, with strain included as a random factor.

We also used a general linear mixed model to evaluate possible factors contributing to differences in survival rates and the probabilities of whole litter loss. We modeled survival rates and the probability of whole litter loss as a function of dam age, the number of previous births, the presence of overlapping litters, and in cases where an overlapping litter was present, the number of days litters overlapped and the number of overlapping siblings. Strain was included as a random factor in the model. In addition, we included litter size as an additional predictor for the survival rate model and weighted the whole litter loss rate by the litter size.

We determined whether there were significant relationships between trio-pair divergence with the female reproductive condition and male sperm quality. We accessed body weights from the McMullan1 dataset and sperm quality data from the Shorter4 dataset, both housed in the Mouse Phenome Database (RRID:SCR_003212).^62,63^ Further, we determined whether there were significant relationships between the trio-pair trait divergence and the average strain reproductive success metrics for overall strain average litter size, interbirth interval, and age at first birth. Finally, we calculated the Spearman correlations to determine the magnitude and direction of the relationship between these traits and the trio-pair trait divergence.

---

**Supplementary Table 1.** The heritabilities of each trait were calculated independently for each breeding configuration as well as for 1,000 replicates of trio-pair divergence values (mean ± standard deviation with minimum and maximum values)

|  | Trio | Pair | Divergence |
| --- | --- | --- | --- |
| Litters per Cross | 0.27 | 0.20 | 0.137 ± 0.009 (0.108, 0.167) |
| Pups per Cross | 0.35 | 0.21 | 0.165 ± 0.010 (0.132, 0.207) |
| Whole Litter Loss Rate | 0.21 | 0.11 | 0.084 ± 0.008 (0.061, 0.114) |
| Interbirth Interval | 0.05 | 0.02 | 0.130 ± 0.014 (0.097, 0.185) |
| Litter Size at Birth | 0.34 | 0.30 | 0.094 ± 0.008 (0.074, 0.124) |
| Litter Size at Wean | 0.34 | 0.26 | 0.115 ± 0.009 (0.089, 0.150) |
| Survival to Weaning | 0.22 | 0.17 | 0.098 ± 0.008 (0.076, 0.125) |

**Supplementary Figure 1.**
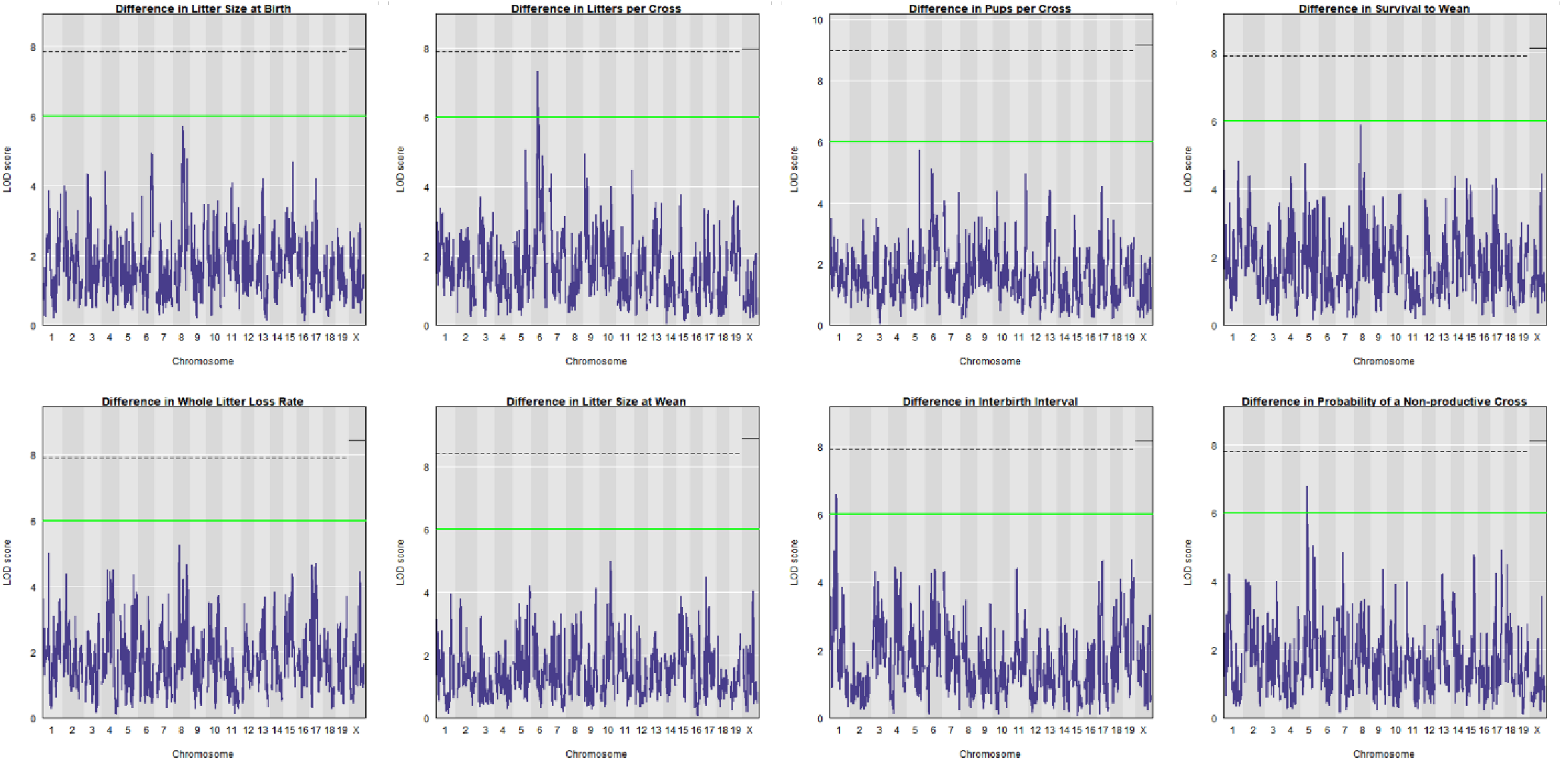
QTL mapping results for the trio-pair differential in various reproductive fitness traits. A LOD score of six is indicated with a green horizontal line, with suggestive peaks crossing that line present for the trio-pair difference in interbirth interval, liters per cross, and the probability of a non-productive cross. The black dotted and solid lines indicate the permutation-based LOD score cutoff of *P*<0.05 on the autosomes and sex chromosomes, respectively.

## Acknowledgments

The authors gratefully acknowledge Racheal Wallace for conserving cage cards from all CC mating units, enabling the compilation of comprehensive breeding records from this mouse population. We also thank Franky Barradale for support in digitizing the cage card records. We are grateful to Uma Arora, Alexandra G. Duffy, Dr. Rebecca E. Forkner, and Kimberly Heath-Borrero providing valuable feedback on manuscript drafts, and the members of the Dumont laboratories for additional helpful discussions and feedback.

## Contributions

A.G. and B.D. contributed to study conceptualization and design. A.G. led the formal data analysis. B.D. led the data collection and transcription effort, assisted by A.G.. A.G. led the writing of the manuscript, supported by B.D.. B.D. contributed to editing and review of the manuscript.

## References

1. West-Eberhard, M. J. Alternative adaptations, speciation, and phylogeny (A Review). Proc Natl Acad Sci U S A 83, 1388–1392 (1986).

2. Taborsky, M., Oliveira, R. F. & Brockmann, H. J. The evolution of alternative reproductive tactics: concepts and questions. in Alternative Reproductive Tactics (eds. Oliveira, R. F., Taborsky, M. & Brockmann, H. J.) 1–22 (Cambridge University Press, 2008). doi:10.1017/CBO9780511542602.002.

3. Taborsky, M. & Brockmann, H. J. Alternative reproductive tactics and life history phenotypes. in Animal Behaviour: Evolution and Mechanisms (ed. Kappeler, P.) 537–586 (Springer Berlin Heidelberg, 2010). doi:10.1007/978-3-642-02624-9_18.

4. Oliveira, R. F., Taborsky, M. & Brockmann, H. J. Alternative Reproductive Tactics: An Integrative Approach. (Cambridge University Press, 2008).

5. Riedman, M. L. The Evolution of Alloparental Care and Adoption in Mammals and Birds. The Quarterly Review of Biology 57, 405–435 (1982).

6. Hrdy, S. B. Mothers and Others: The Evolutionary Origins of Mutual Understanding. (Belknap Press, 2011).

7. Lahdenperä, M., Lummaa, V., Helle, S., Tremblay, M. & Russell, A. F. Fitness benefits of prolonged post-reproductive lifespan in women. Nature 428, 178–181 (2004).

8. Russell, A. F. & Lummaa, V. Maternal effects in cooperative breeders: from hymenopterans to humans. Philos Trans R Soc Lond B Biol Sci 364, 1143–1167 (2009).

9. Burkart, J. M., Hrdy, S. B. & Van Schaik, C. P. Cooperative breeding and human cognitive evolution. Evolutionary Anthropology: Issues, News, and Reviews 18, 175–186 (2009).

10. Kenkel, W. M., Perkeybile, A. M. & Carter, C. S. The neurobiological causes and effects of alloparenting. Developmental Neurobiology 77, 214–232 (2017).

11. Sayler, A. & Salmon, M. An Ethological Analysis of Communal Nursing By the House Mouse (Mus Musculus). Behaviour 40, 62–84 (1971).

12. König, B. Components of lifetime reproductive success in communally and solitarily nursing house mice — a laboratory study. Behav Ecol Sociobiol 34, 275–283 (1994).

13. Manning, C. J., Dewsbury, D. A., Wakeland, E. K. & Potts, W. K. Communal nesting and communal nursing in house mice, Mus musculus domesticus. Animal Behaviour 50, 741– 751 (1995).

14. Dobson, F. S. & Baudoin, C. Experimental tests of spatial association and kinship in monogamous mice (Mus spicilegus) and polygynous mice (Mus musculus domesticus). Can. J. Zool. 80, 980–986 (2002).

15. König, B., Lindholm, A. K. & Pialek, J. The complex social environment of female house mice (Mus domesticus). in Evolution of the House Mouse (eds. Macholan, M., Baird, S. J. E. & Munclinger, P.) 114–134 (Cambridge University Press, 2012). doi:10.1017/CBO9781139044547.007.

16. Palanza, P., Della Seta, D., Ferrari, P. F. & Parmigiani, S. Female competition in wild house mice depends upon timing of female/male settlement and kinship between females. Animal Behaviour 69, 1259–1271 (2005).

17. Ferrari, M., Lindholm, A. K. & König, B. The risk of exploitation during communal nursing in house mice, Mus musculus domesticus. Animal Behaviour 110, 133–143 (2015).

18. Schmidt, J. et al. Reproductive asynchrony and infanticide in house mice breeding communally. Animal Behaviour 101, 201–211 (2015).

19. Garner, J. P., Gaskill, B. N. & Pritchett-Corning, K. R. Two of a Kind or a Full House? Reproductive Suppression and Alloparenting in Laboratory Mice. PLOS ONE 11, e0154966 (2016).

20. Weidt, A., Lindholm, A. K. & König, B. Communal nursing in wild house mice is not a byproduct of group living: Females choose. Naturwissenschaften 101, 73–76 (2014).

21. Auclair, Y., König, B. & Lindholm, A. K. Socially mediated polyandry: a new benefit of communal nesting in mammals. Behavioral Ecology 25, 1467–1473 (2014).

22. Thonhauser, K. E., Raveh, S., Hettyey, A., Beissmann, H. & Penn, D. J. Why do female mice mate with multiple males? Behav Ecol Sociobiol 67, 1961–1970 (2013).

23. Ferrari, M., Lindholm, A. K. & König, B. Fitness Consequences of Female Alternative Reproductive Tactics in House Mice (Mus musculus domesticus). The American Naturalist 193, 106–124 (2019).

24. Hurst, J. L. Behavioural variation in wild house mice Mus domesticus Rutty: A quantitative assessment of female social organization. Animal Behaviour 35, 1846–1857 (1987).

25. König, B., Riester, J. & Markl, H. Maternal care in house mice (Mus musculus): II. The energy cost of lactation as a function of litter size. Journal of Zoology 216, 195–210 (2009).

26. Ferrari, M., Lindholm, A. K., Ozgul, A., Oli, M. K. & König, B. Cooperation by necessity: condition- and density-dependent reproductive tactics of female house mice. Commun Biol 5, 1–10 (2022).

27. Sinervo, B., Chaine, A. S. & Miles, D. B. Social Games and Genic Selection Drive Mammalian Mating System Evolution and Speciation. The American Naturalist (2020) doi:10.1086/706810.

28. Patris, B. & Baudoin, C. Female sexual preferences differ inMus spicilegusandMus musculus domesticus: the role of familiarization and sexual experience. Animal Behaviour 56, 1465–1470 (1998).

29. Ambaryan, A. V., Voznessenskaya, V. V. & Kotenkova, E. V. Mating behavior differences in monogamous and polygamous sympatric closely related species Mus musculus and Mus spicilegus and their role in behavioral precopulatory isolation. Rus.J.Theriol. 18, 67–79 (2019).

30. Green, T. S. O. T. J. L. E. L. Biology of the Laboratory Mouse. (McGraw Hill, 1966).

31. Beck, J. A. et al. Genealogies of mouse inbred strains. Nat Genet 24, 23–25 (2000).

32. Berry, M. L. & Linder, C. C. Chapter 4 - Breeding Systems: Considerations, Genetic Fundamentals, Genetic Background, and Strain Types. in The Mouse in Biomedical Research (Second Edition) (eds. Fox, J. G.et al.) 53–78 (Academic Press, 2007). doi:10.1016/B978-012369454-6/50016-9.

33. Wasson, K. Retrospective Analysis of Reproductive Performance of Pair-bred Compared with Trio-bred Mice. Journal of the American Association for Laboratory Animal Science 56, 190–193 (2017).

34. Heiderstadt, K. M. & Blizard, D. A. Increased Juvenile and Adult Body Weights in BALB/cByJ Mice Reared in a Communal Nest. Journal of the American Association for Laboratory Animal Science 50, 484–487 (2011).

35. Chatkupt, T. T., Libal, N. L., Mader, S. L., Murphy, S. J. & Saunders, K. E. Effect of Continuous Trio Breeding Compared with Continuous Pair Breeding in ‘Shoebox’ Caging on Measures of Reproductive Performance in Estrogen Receptor Knockout Mice. Journal of the American Association for Laboratory Animal Science 57, 328–334 (2018).

36. Sayler, A. & Salmon, M. Communal Nursing in Mice: Influence of Multiple Mothers on the Growth of the Young. Science 164, 1309–1310 (1969).

37. Heiderstadt, K. M., Vandenbergh, D. J., Gyekis, J. P. & Blizard, D. A. Communal Nesting Increases Pup Growth But Has Limited Effects on Adult Behavior and Neurophysiology in Inbred Mice. Journal of the American Association for Laboratory Animal Science 53, 152– 160 (2014).

38. Morello, G. M. et al. High laboratory mouse pre-weaning mortality associated with litter overlap, advanced dam age, small and large litters. PLOS ONE 15, e0236290 (2020).

39. Estep, D. Q., Lanier, D. L. & Dewsbury, D. A. Copulatory behavior and nest building behavior of wild house mice (Mus musculus). Anim Learn Behav 3, 329–336 (1975).

40. Poley, W. Emotionality related to maternal cannibalism in BALB and C57BL mice. Animal Learning & Behavior 2, 241–244 (1974).

41. Weber, E. M., Hultgren, J., Algers, B. & Olsson, I. A. S. Do Laboratory Mouse Females that Lose Their Litters Behave Differently around Parturition? PLOS ONE 11, e0161238 (2016).

42. Mathot, K. J. & Giraldeau, L.-A. Within-group relatedness can lead to higher levels of exploitation: a model and empirical test. Behavioral Ecology 21, 843–850 (2010).

43. Keele, G. R., Crouse, W. L., Kelada, S. N. P. & Valdar, W. Determinants of QTL Mapping Power in the Realized Collaborative Cross. G3 Genes|Genomes|Genetics 9, 1707–1727 (2019).

44. Tarín, J. J. et al. Delayed Motherhood Decreases Life Expectancy of Mouse Offspring1. Biology of Reproduction 72, 1336–1343 (2005).

45. Brajon, S. et al. Social environment as a cause of litter loss in laboratory mouse: A behavioural study. Applied Animal Behaviour Science 218, 104827 (2019).

46. Martínez-Gómez, M., Juárez, M., Distel, H. & Hudson, R. Overlapping litters and reproductive performance in the domestic rabbit. Physiology & Behavior 82, 629–636 (2004).

47. Alsina-Llanes, M., De Brun, V. & Olazábal, D. E. Development and expression of maternal behavior in naïve female C57BL/6 mice. Developmental Psychobiology 57, 189–200 (2015).

48. König, B. Fitness effects of communal rearing in house mice: the role of relatedness versus familiarity. Animal Behaviour 48, 1449–1457 (1994).

49. Rusu, A. S., König, B. & Krackow, S. Pre-reproductive alliance formation in female wild house mice (Mus domesticus): the effects of familiarity and age disparity. acta ethol 6, 53– 58 (2004).

50. Sear, R. & Mace, R. Who keeps children alive? A review of the effects of kin on child survival. Evolution and Human Behavior 29, 1–18 (2008).

51. Haines, B. A., Barradale, F. & Dumont, B. L. Patterns and mechanisms of sex ratio distortion in the Collaborative Cross mouse mapping population. Genetics 219, iyab136 (2021).

52. Churchill, G. A. et al. The Collaborative Cross, a community resource for the genetic analysis of complex traits. Nature Genetics 36, 1133–1137 (2004).

53. Chesler, E. J. et al. The Collaborative Cross at Oak Ridge National Laboratory: developing a powerful resource for systems genetics. Mamm Genome 19, 382–389 (2008).

54. Roberts, A., Pardo-Manuel de Villena, F., Wang, W., McMillan, L. & Threadgill, D. W. The polymorphism architecture of mouse genetic resources elucidated using genome-wide resequencing data: implications for QTL discovery and systems genetics. Mamm Genome 18, 473–481 (2007).

55. Yang, H. et al. Subspecific origin and haplotype diversity in the laboratory mouse. Nat Genet 43, 648–655 (2011).

56. Saul, M. C., Philip, V. M., Reinholdt, L. G. & Chesler, E. J. High-Diversity Mouse Populations for Complex Traits. Trends in Genetics 35, 501–514 (2019).

57. Wahlsten, D., Metten, P. & Crabbe, J. C. A rating scale for wildness and ease of handling laboratory mice: results for 21 inbred strains tested in two laboratories. Genes, Brain and Behavior 2, 71–79 (2003).

58. Chesler, E. J. Out of the bottleneck: the Diversity Outcross and Collaborative Cross mouse populations in behavioral genetics research. Mamm Genome 25, 3–11 (2014).

59. Rutledge, H. et al. Genetic Regulation of Zfp30, CXCL1, and Neutrophilic Inflammation in Murine Lung. Genetics 198, 735–745 (2014).

60. Broman, K. W. et al. R/qtl2: Software for Mapping Quantitative Trait Loci with High-Dimensional Data and Multiparent Populations. Genetics 211, 495–502 (2019).

61. Churchill, G. A. & Doerge, R. W. Empirical threshold values for quantitative trait mapping. Genetics 138, 963–971 (1994).

62. Bogue, M. A., Churchill, G. A. & Chesler, E. J. Collaborative Cross and Diversity Outbred data resources in the Mouse Phenome Database. Mamm Genome 26, 511–520 (2015).

63. Bogue, M. A. et al. Mouse Phenome Database: a data repository and analysis suite for curated primary mouse phenotype data. Nucleic Acids Res 48, D716–D723 (2020).

